# Ten Genetic Loci Identified for Milk Yield, Fat, and Protein in Holstein Cattle

**DOI:** 10.1101/2020.06.17.158386

**Authors:** Liyuan Liu, Jinghang Zhou, Chunpeng James Chen, Juan Zhang, Wan Wen, Jia Tian, Zhiwu Zhang, Yaling Gu

## Abstract

High-yield and high-quality of milk are the primary goals of dairy production. Understanding the genetic architecture underlying these milk production traits is beneficial so that genetic variants can be targeted toward the genetic improvement. In this study, we measured five milk production traits in Holstein cattle population from China. These traits included milk yield, protein yield, fat yields; fat percentage and protein percentages. We used the estimated breeding values as dependent variables to conduct the genome-wide association studies (GWAS). Breeding values were estimated through pedigree relationships by using a mixed linear model for individuals with and without phenotypic data. Genotyping was carried out on the individuals with phenotypes by using the Illumina BovineSNP150 BeadChip. The association analyses were conducted by using the Fixed and random model Circulating Probability Unification (FarmCPU) method. A total of ten SNPs was detected above the genome-wide significant threshold, including six located in previously reported QTL regions. We found eight candidate genes within distances of 120 kb upstream or downstream to the associated SNPs. The most significant SNP is on *DGAT1* gene affecting milk fat and protein percentage. These genetic variants and candidate genes would be valuable resources to enhance dairy cattle breeding.

## Introduction

Traditional dairy breeding mainly depends on phenotypic and pedigree information that is expensive, labor-intensive, and time-consuming. Dairy cows have a long generation interval and give birth only once a year. With the advances in molecular genetic techniques, genomic selection (GS) has been widely used in plant and animal breeding. It was demonstrated that implementation of genomic selection can increase genetic gain in dairy cattle [1]. Many statistical methods for GS have been proposed, and the method of integrating Genome-wide association studies (GWAS) and GS into one step has been developed, which account the genetic architecture of interest traits [2]. The conventional genomic best linear unbiased prediction (GBLUP) considering all markers that have the same effects, so thus ignores the markers that have large effect on target traits.

Research showed that the prediction accuracy was higher than that use of conventional GBLUP when fitting the most significant marker in the prediction model [3]. Also Resende et al suggested that the prediction accuracy reached maximum when constructed the genomic relationship matrix using causative quantities trait nucleotides (QTNs) [4]. Incorporating the public most significant markers from GWAS showed that it can improve the prediction accuracy in dairy cattle [5,6]. And the prediction accuracy were higher when accounting for variants from GWAS results [7]. Therefore, GWAS are very helpful for further GS. Since GWAS have proven to be a powerful method for identifying potential genetic variants, especially single-nucleotide polymorphisms (SNPs) associated with complex traits in humans and animals [8–10].

Milk production and milk composition are the most important economic traits in dairy industry. A large number of association studies reveals numerous QTLs for milk-related traits in dairy cattle population around the world over the past 20 years (CatttleQTLdb: https://www.animalgenome.org/cgi-bin/QTLdb/BT/index), and many researchers conducted meta-analysis based on GWAS results for milk-related traits in different cattle breeds [7,11,12]. Although early study revealed 105 SNPs associated with milk production traits in Chinese Holstein population, but they used a lower density marker which contains only about 50,000 SNPs [13].

Research showed that high-density genotype could provide markers close to the QTL and help in fine mapping of causative mutation[14]. Also Vanraden reported that high-density marker increased the precision of QTL detection in cattle population[15]. A maize study showed that the GWAS detection efficiency was improved when increased the marker density for husk traits [16]. In addition, several studies reported that the genomic prediction accuracy increased when increased the marker density in cattle breed[17] and shrimp breed[18],Therefore, it is necessary to use dense genotype to identify genetic variation and implementation of genomic prediction.

In this study, we conducted GWAS using Illumina BovineSNP150 BeadChip which contains about 150,000 SNPs. The population was from Holstein cows raised in the Ningxia area of northwest China. Our objectives were to identify new genetic variants associated with five milk traits, including milk yield, fat yield, protein yield, fat percentage, and protein percentage. We expect the newly identified genetic variants and potential candidate genes would become valuable resources for genomic evaluation.

## Material and methods

### Population and phenotypic data

The study population is Holstein cows that were raised on 22 dairy farms in the Ningxia area of China. We used estimated breeding values (EBVs) as phenotypes implement association analysis, Milk yield (MY), fat yield (FY), protein yield (PY), fat percentage (FP), protein percentage (PP) measurements were recorded once a month for each cow after calving and the start of lactation. The milk yield was automatically recorded by the milking system in each farm, the milk components are tested by the Dairy Herd Improvement lab at Animal Husbandry of Extension Station in Ningxia, using spectrometers. FY was calculated as (FP*MY)/100; PY was calculated as (PP*MY)/100. The distribution of phenotypes and correlations between the different phenotypic traits are illustrated in **Figures** S1.

### Estimated breeding values

The DMU package (Derivative-free approach to MUltivariate analysis) [19] was used to estimate breeding values using Random Regression Test-Day Model [20,21]. Totally about 452,920 test-day records from 61,600 cows spanning a 9-year period (2011-2019) at their first lactation stage, all recorded data reflect the first lactation for each cow. We considered herd-test-day and calving year-season as fixed effects, calving month-age as a fixed regression effect, and individual additive effect and permanent environment effect as random regression effects. Fixed regression using a fourth order Legendre polynomial. The model equation is as follows:

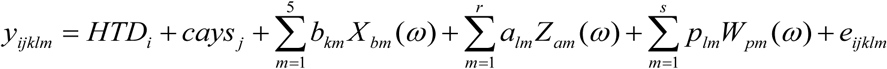

Where *y*_*ijklm*_ is the test-day records m of cow ; *HTD*_*i*_ is the fixed effect of the i th herd-test day; *cays*_*j*_ is the fixed effect of the j th combined of calving years and season; *b*_*km*_ is the m th fixed regression coefficient of the k th calving year-season, *X*_*bm*_ is the Legendre polynomial m th covariable; *a*_*lm*_ is the m th random regression coefficient under additive genetic effects, corresponds to the l th cow in the pedigree; *p*_*lm*_ is the m th random regression coefficient under the permanent environment genetic effects, correspond to the l th cow in the data file; *Z*_*am*_ and *W*_*pm*_ are the m th covariance elements for additive effects and permanent environment effects; *ω* is the days of lactation after standardization; and *e*_*ijklm*_ is the residual effects. A heatmap of estimated breeding values for milk production traits is illustrated in **Figure** S2.

### Genotypic data

Blood samples from the 1,220 cows were collected by cattle farm staff in this study. DNA was extracted and Genotyping was carried out by Compass Biotechnology (http://www.kangpusen.com/) using the Illumina BovineSNP150 BeadChip. We used genome reference the Bos_taurus_UMD_3.1. In total, there were 124,743 variants for the association analysis after conducted quality control by Plink software [22]. Markers were removed if (1) the call rate of an individual genotype was less than 95%, (2) the call rate of a single SNP genotype was less than 90%, and (3) if the MAF of a SNP was less than 0.05 and deviated from Hardy-Weinberg equilibrium (p < 1.0e-6). We calculated marker intervals and linkage disequilibrium (LD) to estimate R square for all markers and plotted the marker distribution in **Figure** 1.

**Figure 1.**
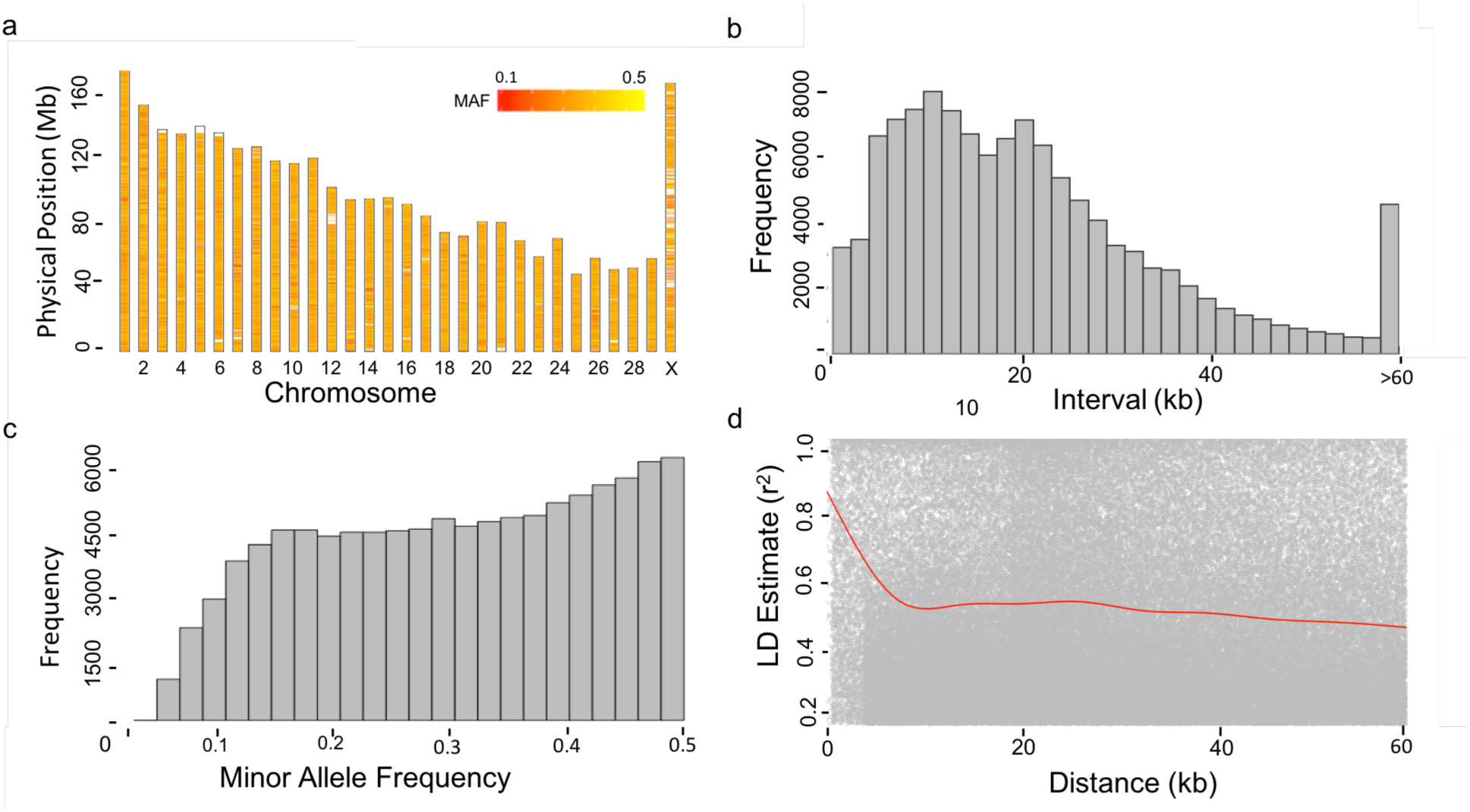
Properties of Single Nucleotide Polymorphisms (SNPs). A total of 1,220 cows were genotyped by the Illumina Bovine 150 k BeadChip. After conducting quality control on both minor allele frequency (MAF, above 5%) and missing rate (< 10%), 1,220 individuals and 124,743 SNP remained. The distribution of the filtered SNPs is displayed over the 30 bovine chromosomes except for Y chromosome (a). The MAFs of SNPs were re-calculated after the filtering and were displayed by a heat map. Consequently, there SNPs with MAF < 5% remained, as demonstrated by the histogram (c). The density of SNPs is displayed by the frequency of the distance between adjacent SNPs (b). The distances over 60 kb clustered into one group. The maximum distance was 100.06 kb. Pairwise Linkage Disequilibrium (LD) was calculated as the R square for SNPs within the 100 kb window. The decay of LD over distance (red line) is displayed by the pairwise LD and moving average (d).

### Principle component analysis

*P*rinciple component analysis (PCA) was conducted on 1,220 cows genotyped with 124,743 markers covering the whole genome to detect the population structure. The 1,220 cows are the progeny of the frozen semen imported from multiple countries. The of origins of the sires were plotted against the principal components to demonstrate how the population structure was formed.

### Association analysis

We performed the single-SNP analysis for the markers using the FarmCPU (Fixed and random model Circuitous Probability Unification) [23]. The genome wise threshold corresponding type I error of 1% was 4.0e-07 after Bonferroni multiple test correction (5%/124,743). FarmCPU can consider a population structure matrix as a covariate variable to correct false positive results caused by population stratification [23]. Considering the stratification in the dairy cow population, we fit the first three principal components as covariate variables in the GWAS models.

### Annotation of candidate gene and pathway analysis

The genome reference Bos_taurus_UMD_3.1 was used to search candidate gene. The candidate genes were found within distances of 120 kb upstream or downstream to the associated SNPs. The online website “https://oct2018.archive.ensembl.org/Bos_taurus/Info/Index”, “https://www.ncbi.nlm.nih.gov/gene/”, “https://www.genome.jp/kegg/pathway.html” were used for functional analysis and pathway analysis of the candidate gene by GWAS.

## Results

### Phenotypic and estimated genetic parameters

Phenotype distributions and correlations among phenotypic traits, estimated breeding values, and residuals are shown in the supplementary material (**Figure** S1). There were strong positive phenotypic correlations between “yield” type of traits, including MY, PY, and FY. Their phenotypic correlations were 0.90 (MY and PY), 0.70 (MY and FY), and 0.74 (PY and FY).

We also found strong positive genetic correlations between MY and PY (*r*_*g*_ = 0.92), MY and FY (*r*_*g*_ = 0.84), and PY and FY (*r*_*g*_ = 0.88). In contrast, there were weak negative phenotypic and genetic correlations between MY and FP (*r*_*p*_ = −0.15, *r*_*g*_ = −0.32), MY and PP (*r*_*p*_ = −0.20, *r*_*g*_ = − 0.44).

In this study, we used EBVs as the dependent variables for GWAS, **Figure** S2 showed that the heatmap of EBVs for five milk traits. A test-day model to estimate the heritability for each trait and breeding values for individuals. The heritability estimates for MY, FY, PY, FP and PP were 0.12, 0.18, 0.27, 0.30 and 0.32, respectively (**Table** 1).

**Table 1.** Variance components and heritability of milk traits*.

| Variance component | MY | FP | FY | PP | PY |
| --- | --- | --- | --- | --- | --- |
| Genetic | 1592.62 | 4.76 | 1.48 | 1.65 | 1.20 |
| Permanent environmental | 4267.93 | 10.58 | 5.46 | 3.45 | 3.96 |
| Residual | 7413.63 | 0.49 | 0.08 | 0.07 | 0.03 |
| $h^2$ | 0.12 | 0.30 | 0.21 | 0.32 | 0.23 |

\*MY, milk yield; FP, fat percentage; FY, fat yield; PP, protein percentage, PY, protein yield.

### Marker information

We conducted GWAS analyses with 1,220 Holstein dairy cows and 124,743 markers after quality control (QC). Markers covered all 29 autosomes plus the X sex chromosome (**Figure 1a**). After the QC filtering, we re-calculated the minor allele frequency (MAF) for all SNPs. The minimum MAF was 3.8%. There were only 0.1% of markers with MAF below 5% (**Figure 1c**). Marker density was high. Majority of markers (56%) are within 20 kb distances to their adjacent markers (**Figure 1b**). Within such distance, the LD was strong (average R^2^ = 0.56) (**Figure 1d**).

### Population structure

To determine the level of population stratification, we plotted the population structure by principal component analysis (PCA). The population stratified into two unevenly sized groups (**Figure 2**). We also made scatter plot of bull’s country source (**Figure** S3). To adjust for the population stratification, we fit the first three principal components (PCs) as covariate variables in the association analysis. We also made a scatter plot between the first three PCs and the five traits, there are weak correlation observed between the PCs and these phenotypes (**Figure** S4).

**Figure 2.**
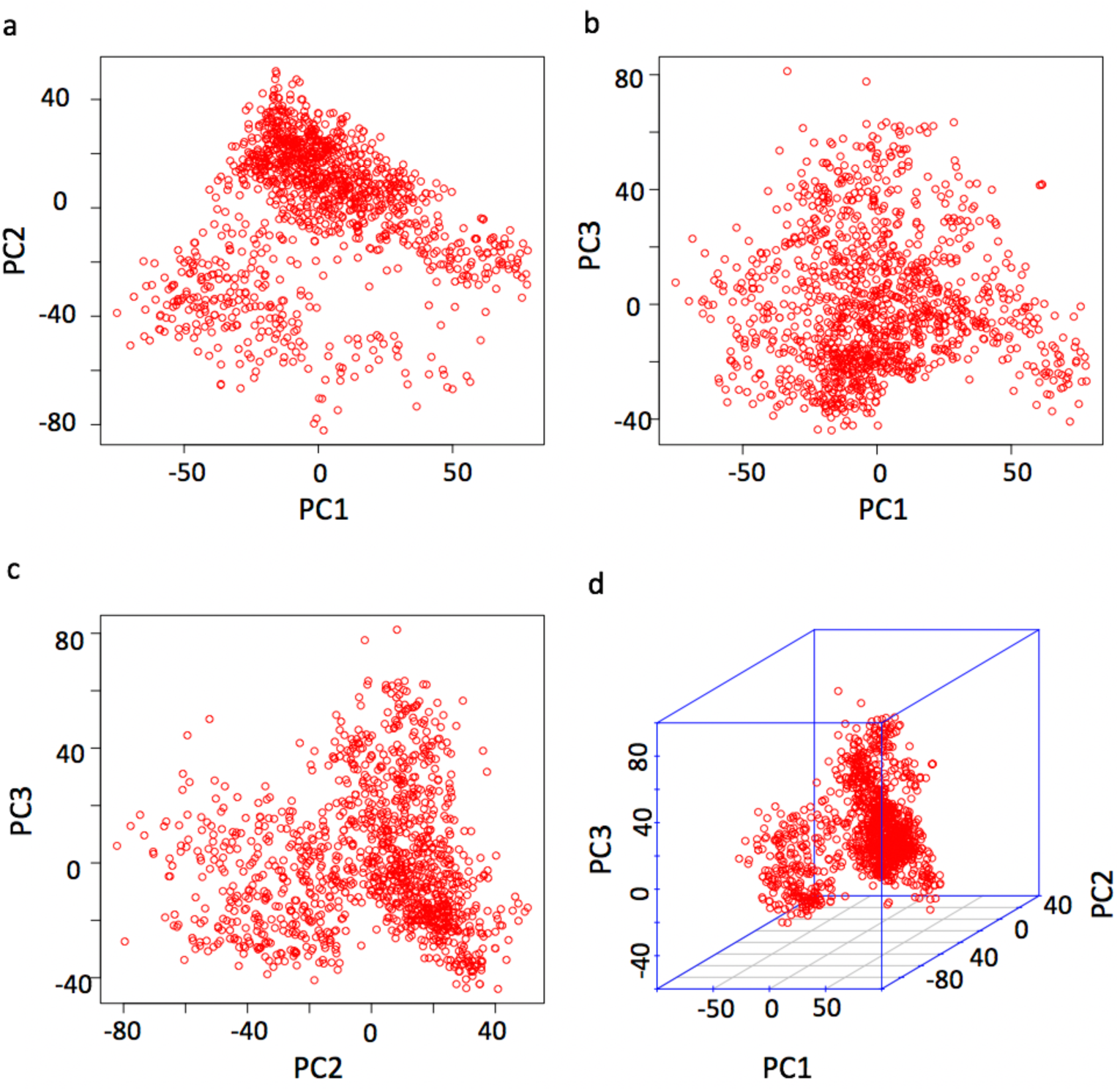
Population structure demonstrated by principal component analysis. Principal Component Analysis (PCA) was conducted with the 124,743 SNPs for the 1,220 cows. The population structure is demonstrated by the pairwise scatter plots (a, b, and c) and the 3D plot (d) of the first three principal components (PCs).

## Results

By drawing the Quantile-Quantile (QQ) plots, we found that the model for GWAS analysis in this study was reasonable, and the point at the upper right corner also shown that some significant markers were found that associated with four milk traits (**Figure** 3). We used *P* < 4.0e-07 as threshold, which was corresponding to 1% of type I error after Bonferroni multiple test correction. We did not find threshold significant SNPs associated with MY, a total of ten highly significant SNPs associated with four traits (**Table** 2, **Figure** 3). Three SNPs (rs42295213, rs136949224 and rs109421300) associated with FP are located on BTA1, 8 and 14, four SNPs (rs43526055, rs137676276, rs109528658 and rs135780687) associated with FY are located on BTA7, 11, 17, and X, respectively. Three SNPs (rs109875012, rs109421300 and rs108996837) associated with PP are located on BTA5, 14, and 21, respectively. One SNP associated with PY is located on BTA5. Four of these ten significant SNPs are located inside genes *EPHA6* (EPH receptor A6), *SLCO1A2* (solute carrier organic anion transporter family member 1A2), *DGAT1* (diacylglycerol O-acyltransferase 1) *and EP400* (E1A binding protein p400). The SNP (rs109875012) on BTA5 located close to the *ZNF384* (zinc finger protein 384). The SNP (rs10705865) on BTA8 located close to *SCARA5* (scavenger receptor class A member 5) gene. The SNP (rs137676276) on BTA 11 located close to *VIT* (vitrin). The SNP (rs108996837) on BTA 21 located close to *EXOC3L4* (exocyst complex component 3 like 4), and the SNP (rs135780687) on X chromosome located close to *GRPR* (gastrin releasing peptide receptor). The most significant SNP (rs109421300) associated with both FP and PP located in the *DGAT1* gene. Two SNPs (rs137676276, rs108996837) exhibit notably smaller MAFs compared to other SNPs, 0.11 and 0.12, respectively (**Table** 2).

**Figure 3.**
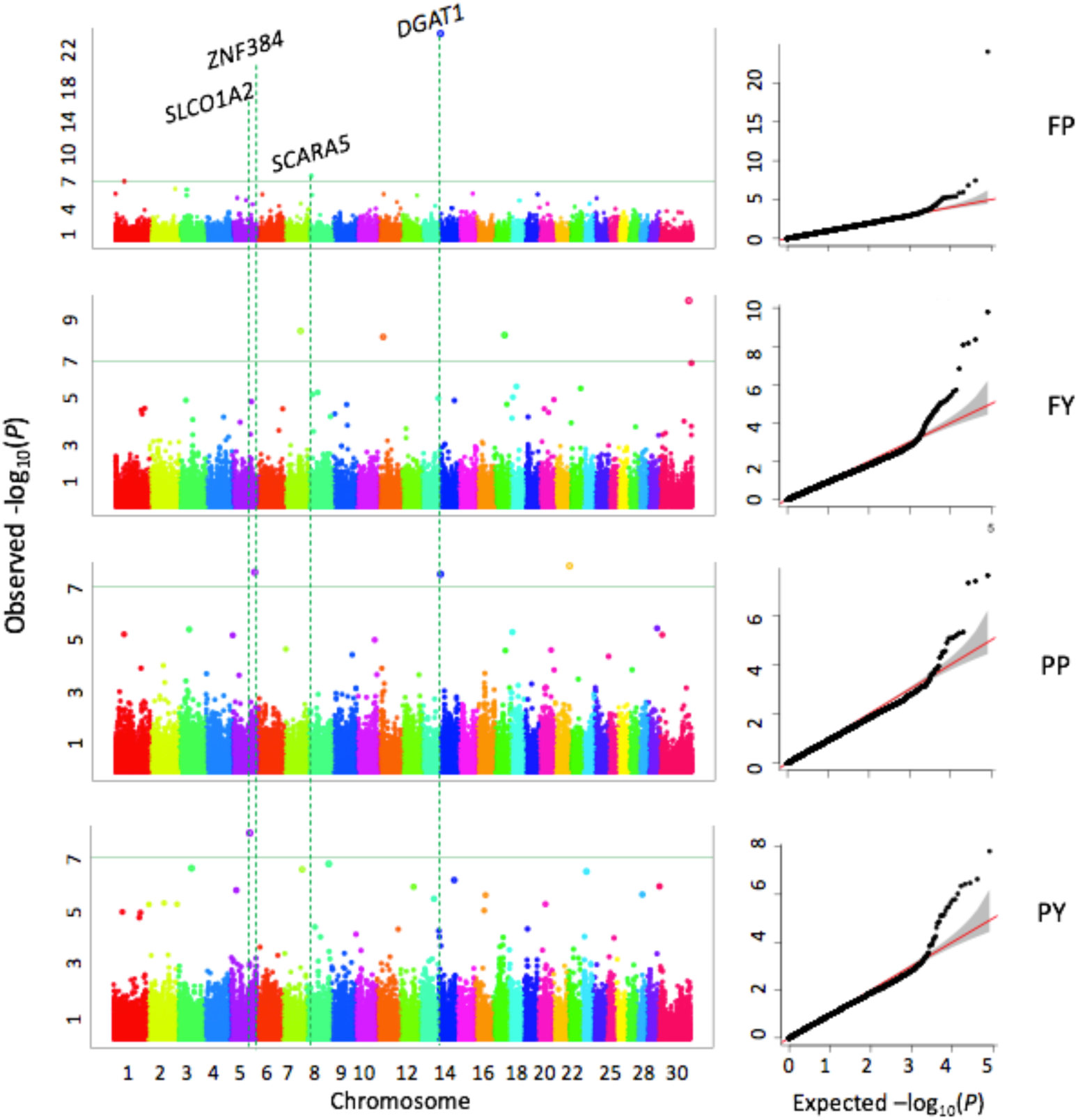
Associations between 124,743 SNPs and milk traits. Milk traits include fat yield (FY), protein yield (PY), fat percentage (FP), protein percentage (PP). The association analyses were conducted by the FarmCPU R package. Manhattan plots display the negative logarithms of the observed *P* values for SNPs across 30 bovine chromosomes (left panel). The green line indicates the Bonferroni multiple test threshold at *P* = 4.0e-07. The Quantile-Quantile (QQ) plots represent the negative logarithms of the expected *P* values (X-axis) and observed *P* values (Y-axis) (right panel).

**Table 2.**
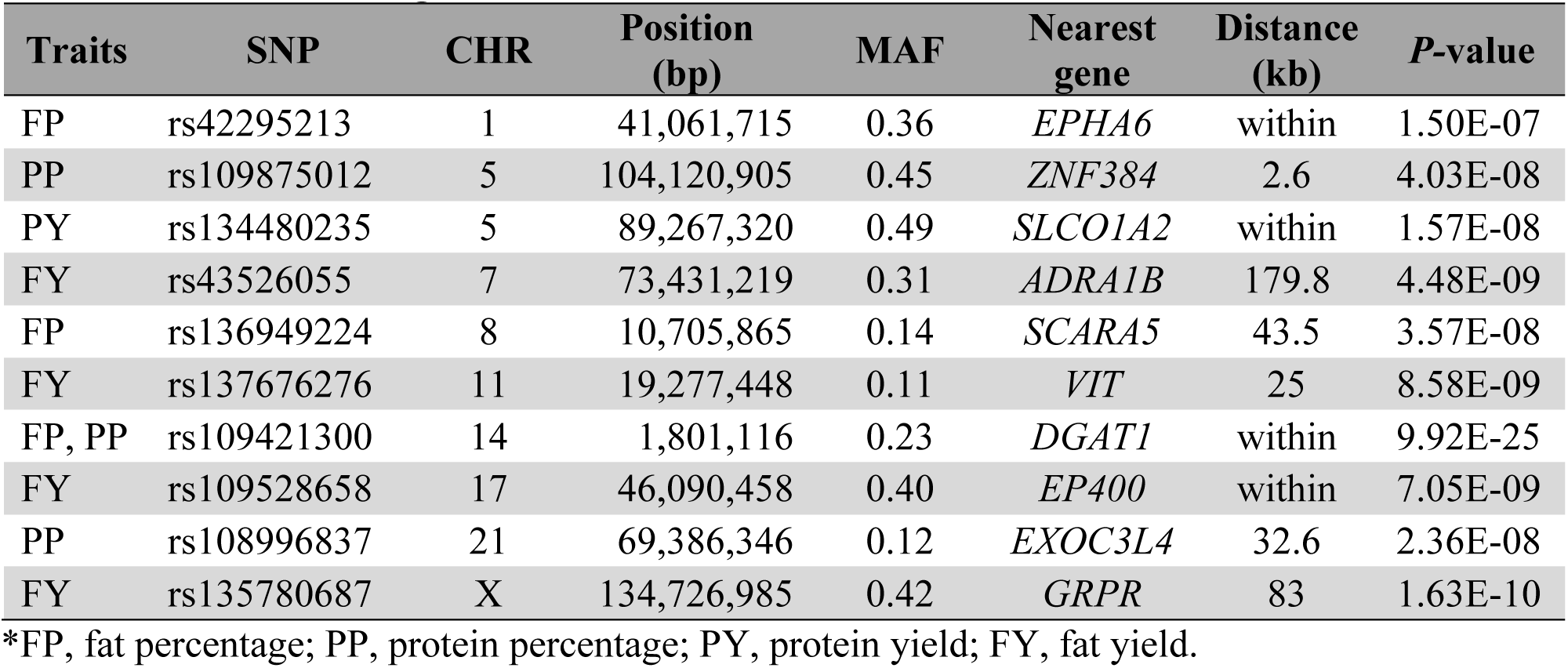
Genome-wide significant SNPs associated with milk traits*.

### Pleiotropic QTLs for milk production traits

We used the SNPs with *P* < 0.0005 to make heatmap to look for markers associated with two or more milk production traits (**Figure** S5). We expected to find pleiotropic QTLs because these milk traits are moderately or highly correlated (**Figure** S1, S2). QTL on BTA1 at 0.4Mb, on BTA2 at 14.8Mb, on BTA2 at 134.3Mb, on BTA3 at 64.4Mb, on BTA11 at 39 Mb, on BTA17 at 29-32.9Mb, on BTA20 at 67.8 Mb, BTA22 at 54.6Mb associated with three milk production traits (MY, PY, and FY). QTL on BTA1 at 40.3-41.1Mb, on BTA6 at 37.7Mb, on BTA14 at 73-77Mb, on BTA17 at 29-32Mb associated with MY and PY. QTL on BTA20 at 24Mb associated with FY and FY.

## Discussion

### Population structure

Population stratification is an important confounding factor due to systematic ancestry differences that can cause false positives in GWAS [24]. Partially in cattle population, not all the registered cattle are the full blood. This point can be explained by looking at the cattle breed registration requirements in different countries. Here we take the Holstein cattle as an example, one of the requirements to register a Holstein cattle in China is that the cattle at least has 87.5% blood of Holstein (Chinese Holstein, GB/T 3157 2008), all the animals with Holstein genetics can be registered in Canadian (https://www.holstein.ca/Public/en/Services/Registration/Registration_Eligibilities) and there are the similar clause in USA (http://www.holsteinusa.com/animal_id/register.html). According to the above standards we can see the common ground to register a Holstein cattle between different countries is that not all the Holstein cattle are the full blood. Even most of the Holstein cattle are pure breed, some of registered cattle could still contains a little other blood in the long-term breeding progress. That’s why the population structure analysis is necessary in this study.

By the principle component analysis, the PCA scatter plot showed that there are two groups in the population in this study; that is, a small number of individuals divided into the other group (**Figure** 2, **Figure** S3). There are probably two reasons caused the deviation of the population structure, the first reason is the semen the cattle farm used are different, as we know most of the farms in this study participated in a dairy breeding project that introduce of some Holstein frozen semen overseas annually from 2013 to 2018. Another reason is that some cows are introduced from different countries there might has some various background.

As we observed the population structure, the principle components were fitted as covariance do association analysis to correcting population stratification. After adjusted PCA factors, there are four SNPs are overlapped with association model not fitted PCs, will discuss that later.

### GWAS for milk production traits

Milk yield and composition are very important economic traits in dairy cattle because good milk production performance can bring greater economic benefits. As early as 1994, a research identified a QTL significantly associated with FY was linked to kappa-casein and a QTL for PY was linked to beta-lactoglobulin [25]. Subsequently, a growing number of studies detected thousands and tens of QTLs through the 30 chromosomes associated with 646 different traits in cattle (Cattle QTLdb). In this study, there were no significant SNPs passed the Bonferroni correction threshold for MY, but ten SNPs were significantly associated with the other four milk traits (FP, FY, PP and PY).

In this study, we found three SNPs on BTA1, 8, 14 were associated with FP, they are within reported QTLs [26–29]. By search the candidate genes within distances of 120 kb upstream or downstream to the associated SNPs, the genome reference is Bos_taurus_UMD_3.1. The SNP on BTA1 located in the *EPHA6* gene, which have functions of ATP binding (GO:0005524), protein binding (GO: 0005515), protein tyrosine kinase activity (GO: 0004713), and has been proposed to participate in Axon guidance pathway (KEGG: bta04360). The SNP on BTA8 located close to *SCARA5* gene, this gene is a member of the scavenger receptor (SR) family, which is broad expression in fat tissue in human individuals, research showed that scavenger receptors involved in lipid accumulation and inflammation [30], some studies reported the gene plays a critical role in progression and metastasis of breast cancer [31], and involved in breast carcinogenesis [32].

*SCARA5* gene may be a potential candidate gene involved in the biosynthesis of milk fat. The SNP on BTA14 associated both with FP and PP, which is in the *DGAT1* gene. As we know, *DGAT1* gene is widely reported associated with milk yield and composition, especially K232A polymorphism affecting on milk fat and protein [27,33,34]. In this study, the SNP (rs109421300) we identified is within an intron of *DGAT1* gene is the most significantly affecting FP (*P* = 9.92e-25), and also affecting PP (*P* = 7.05e-9), and a lot of studies have reported for milk yield and composition in Holstein, Jersey and Holstein-Friesian cattle [35–37]. *DGAT1* gene encodes an multipass transmembrane protein that functions as a key metabolic enzyme, involved in triacylglycerol biosynthesis, and glycerolipid metabolism (KEGG: bta00561), retinol metabolism (KEGG: bta00830), metabolic pathways (KEGG: bta01100) and fat digestion and absorption (KEGG: bta04975). The PP-associated SNP on BTA5 is within previously reported milk fatty acid content QTL [28], this SNP is close to the *ZNF384*, which is encodes a C2H2-type zinc finger protein, it may function as a transcription factor and have DNA binding and metal ion binding functions. Another PP-associated SNP on BTA21. Four SNPs on BTA7, 11, 17 and X chromosome are associated with FY. The SNP on BTA11 at 19Mb located near the VIT, this gene encodes an extracellular matrix (ECM) protein, which has function of glycosaminoglycan binding (GO:0005539), is has been proposed participate in growth plate cartilage chondrocyte morphogenesis pathway (GO:0003429) and nervous system development pathway (GO:0007399). The SNP on BTA17 is within previously reported QTL for milk protein composition [38], and it also located in the *EP400*, this gene has been participate in histone H2A acetylation, histone H4 acetylation, and has the function of ATP binding, DNA binding, protein binding and helicase activity. The FY-associated SNP on chromosome X was located nearby the *GRPR*, this gene is gastrin releasing peptide receptor, gastrin-releasing peptide regulates numerous functions of gastrointestinal and central nervous system, which involved in calcium signaling pathway and neuroactive ligand-receptor interaction. Song reviewed a lot of studies on calcium signaling pathway participated in the effect of the sympathetic never in regulating adipose metabolism [39]. We suspect that *GRPR* gene may affect milk fat through the calcium signaling pathway. Because of the complexity of the biological mechanisms of quantitative traits, the study is only based on SNP data, many studies showed that copy number variation and DNA methylation are also associated with milk performance[40–42], The PY-associated SNP on BTA5 is within previously reported QTL region, which is associated with milk fatty acid content, this SNP is in the *SLCO1A2* gene, which is encoding solute carrier anion transporter family, member 1A2, the gene participate in digestive system, organic anion transport process and bile secretion pathway (KEGG: 04976). Furthermore, when the association model not fitted PCs as covariance, four SNPs are overlapped with model with fitted PCs:rs109875012 (*ZNF384*), rs136949224 (*SCARA5*), rs109421300 (*DGAT1*), and rs109528658 (*EP400*), it is suggested that these significant SNPs could more stable and reliable.

### Correlations among milk traits and pleiotropic QTLs

Estimates of heritability and genetic correlations are essential population genetic parameters in animal breeding research and application of animal breeding programs [43]. Genetic correlation can be useful for indirect selection, selection in different environment and selecting multiple traits simultaneously. We found there were relatively high genetic correlation between milk yield and protein yield, is consistent with the results in Holsteins [44] and the results in Jersey dairy cattle [43]. We expect that there are overlapping region of genetic variation between different traits that are relatively high correlation. A heatmap using *P*-values from GWAS results helped identify pleiotropic QTLs (**Figure** S5). The QTL on BTA2 at 134Mb associated with MY, PY and FY, and also reported associated with the milk composition in another Holstein study [45].

The MY- and PY-associated QTL is adjacent to the *ABCG2* gene on BTA 6, previously studies reported a missense mutations in the *ABCG2* gene is associated with milk yield and composition in Holstein, Braunvieh, and Fleckvieh cattle [11,46], a intron variant affecting milk fatty acids in Chinese Holstein [47], and also reported for body weight, calving ease direct in US cattle breed [48,49]. The QTL on BTA14 in the *DGAT1* gene associated with milk yield, fat and protein percentage. The QTL on BTA17 associated with MY, PY and FY, some study have pointed to QTL on this region related to milk fatty acid and body weight [45,50]. A previous study reported a QTL on BTA 20 is associated with PP and body weight[51,52], in this study, we found this QTL is associated with MY, PY and FY. The QTL on BTA24 is pleiotropic and is associated with MY and FY, Ibeagha-Awemu and Peter reported for milk fatty acid [53], Kiser and Clancey reported for conception rate in Holstein cows [54].

## Conclusions

We performed genome-wide association studies using test-day records for five milk production traits (milk yield, fat yield, protein yield, fat percentage, and protein percentage) in Holstein cows. In total of 10 significantly SNPs associated with fat and protein percentage, fat and protein yield, five SNPs are located within previously reported QTLs. An important top variant in the *DGAT1* gene affecting milk fat and protein, and we also found *SCARA5* and *GRPR* gene may be potential candidate gene involved in the biosynthesis of milk fat. In addition, some pleiotropic QTLs on BTA1, 2, 3, 6, 11, 14, 17, 20, and 24 associated with more than two milk traits, these results could provide some basis for molecular breeding and targets for gene cloning.

## Acknowledgements

This study was partially supported by the High-Level Academic Papers in Ningxia University, the project of High-yield and High-quality Dairy Cattle Breeding (2019NYYZ05), the National Science Foundation (Award # DBI 1661348), and the USDA National Institute of Food and Agriculture Hatch project (1014919). The authors would like to thank the DHI measurement center of Ningxia Animal Husbandry of Extension Station and 21 dairy farms for their phenotypic data. We also thank Dr. Linda R. Klein for valuable writing advice and editing the manuscript.

## Author contributions

Conceived experiment: Zhiwu Zhang and Yaling Gu; Data analyses: Liyuan Liu, Jinghang Zhou, and James Chen; Data collection: Juan Zhang, Jia Tian, and Wan Wen; Wrote manuscript: Liyuan Liu, Jinghang Zhou, James Chen, Zhiwu Zhang, and Yaling Gu

## Ethics Statement

No applicable. This study only did statistical analysis based on the former database which come from the Holstein breeding project in Ningxia province, China. All the authors in this study didn’t participated in any sample collection process. All the farms involved were consent and agree to take part in this research.

