## supplemental for "Ten Genetic Loci Identified for Milk Yield, Fat, and Protein in Holstein Cattle"

### Supplementary material

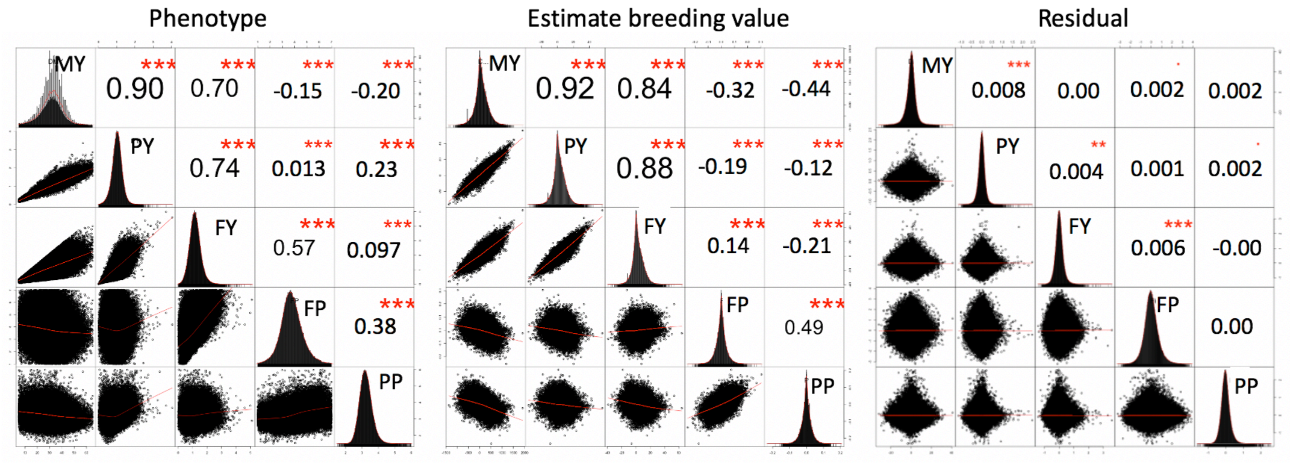


**Figure S1. Phenotype distributions and correlations among milk traits.** The milk traits are demonstrated as phenotypical values (left panel), estimated breeding values (middle panel), and residuals (right panel). The distributions of these values are displayed on the diagonals and the correlations are illustrated as scatter plots off the diagonals. The milk traits include milk yield (MY), fat yield (FY), protein yield (PY), fat percentage (FP), and protein percentage (PP).

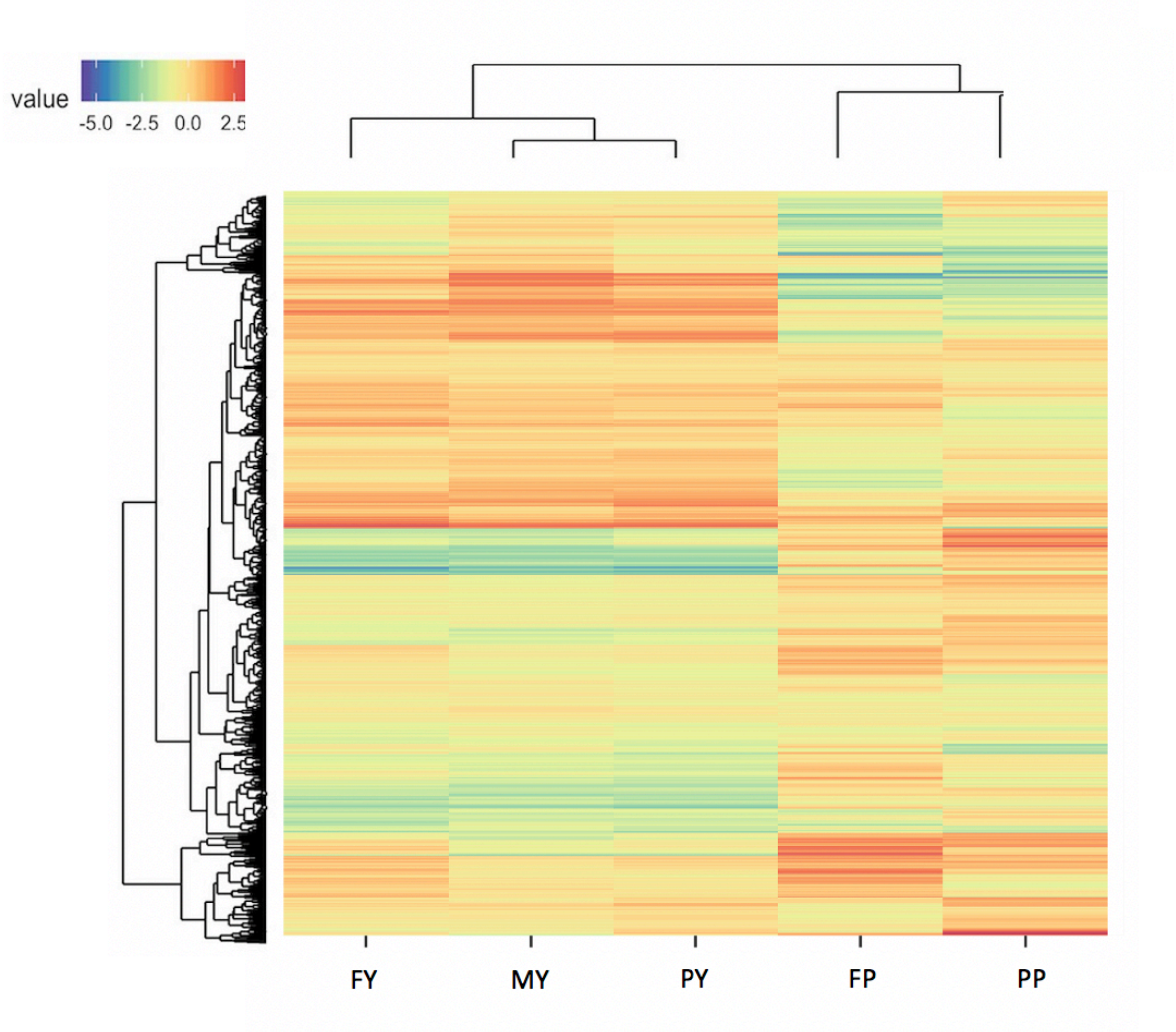


**Figure S2. Heatmap of estimated breeding values for milk traits. Individuals are sorted row wise and traits column wise based on their similarity**. The trait values were standardized and illustrated as heat map with red indicating highest and yellow the lowest. Traits include Fat Yield (FY), Milk Yield (MY), Protein Yield (PY), Protein Percentage (PP), Fat Percentage (FP), and Somatic Cell Score (SCS).

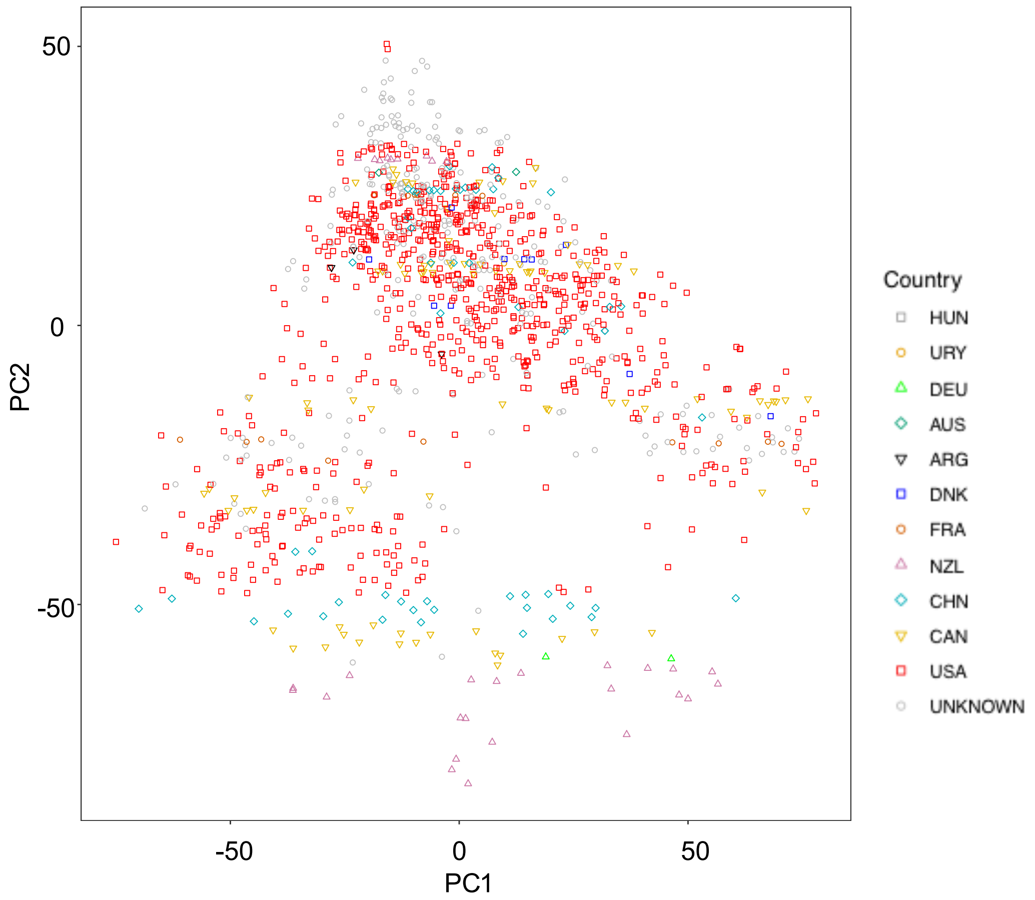


**Figure S3. The relationship between origins and the first two principle components.** The principal components were calculated from all the genetic markers (124,743 SNPs) for the 1,220 cows. Each dot represents a cow with colors and shapes indicating the origins of its sire. HUN, Hungary; URY, Uruguay; DEU, Germany; AUS, Australia; ARG, Argentina; DNK, Denmark; FRA, France; NZL, New Zealand; CHN, China; CAN, Canada; USA, United State of America; UNKNOWN, the country source of the bulls is unknown.

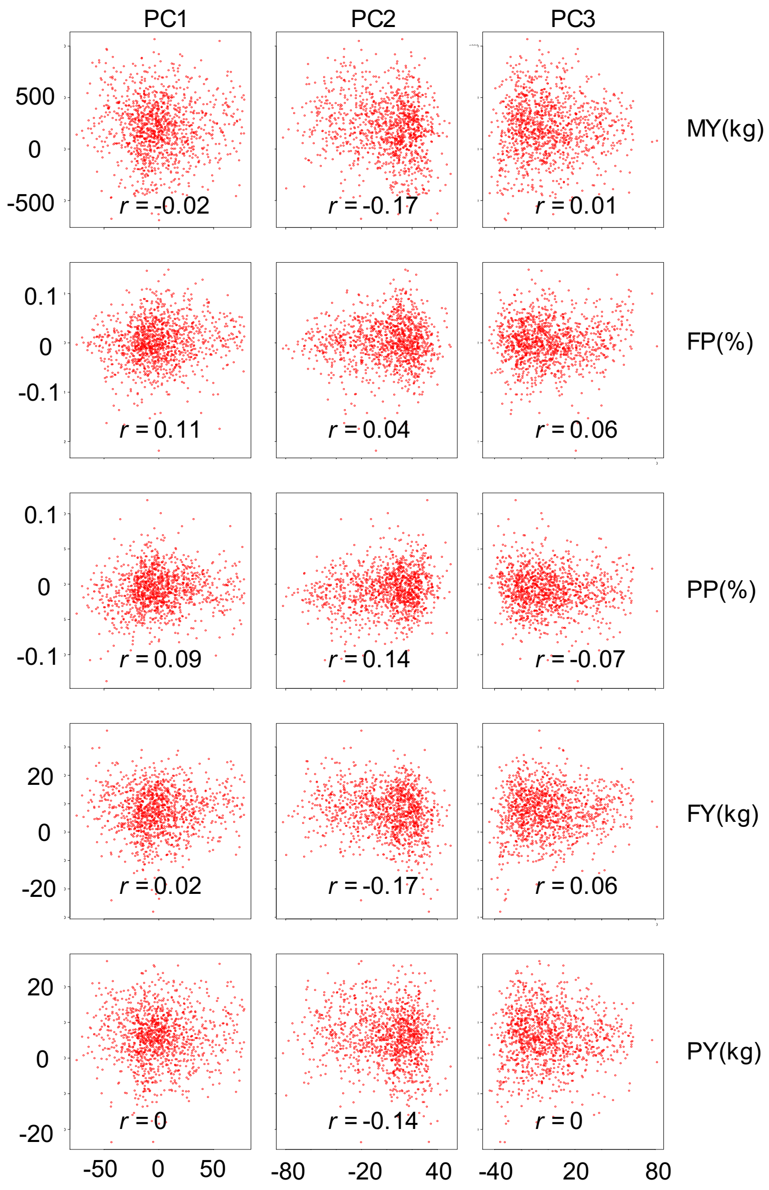


**Figure S4. Correlation between phenotypic values of milk traits and the first three principal components.** The principal components were derived from all genetic markers (124,743 SNPs).

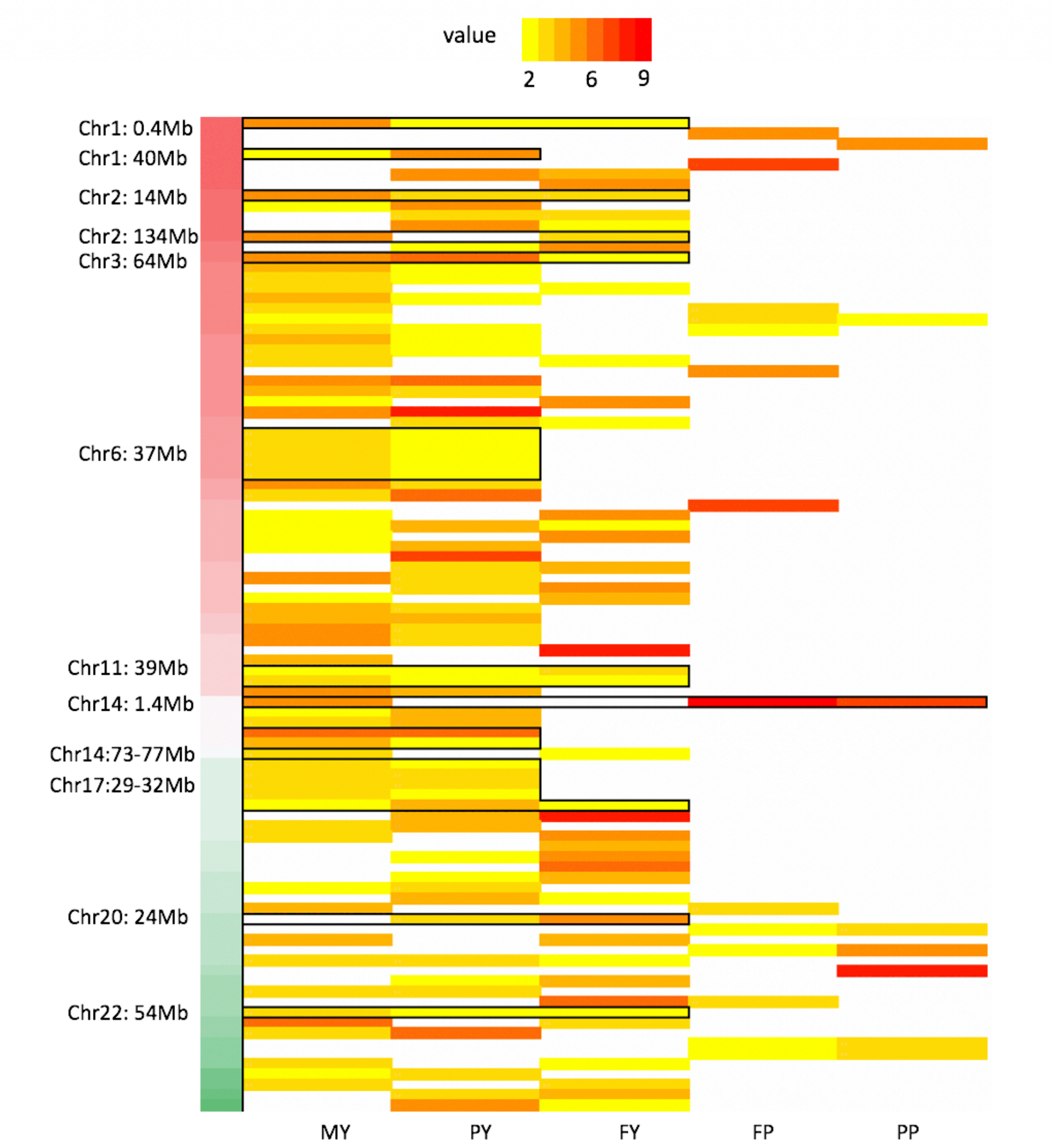


**Figure S5 Heatmap of significant markers associate with five milk traits.** The traits values were standardized and illustrated pleiotropic SNPs. SNPs are associated with more than two traits, one of which have a *P* value less than 10e-04. The traits *P* values illustrated as heat map with red indicating highest and yellow the lowest. The traits include milk yield (MY), fat yield (FY), protein yield (PY), fat percentage (FP), and protein percentage (PP).
